# The hippocampus and dorso-lateral striatum link distinct types of memories through time and space, respectively

**DOI:** 10.1101/2019.12.16.878470

**Authors:** Janina Ferbinteanu

## Abstract

Several decades of research have established that different kinds of memories result from the activity of discrete neural networks. Studying how these networks process information in experiments that target specific types of mnemonic representations has provided deep insights into memory architecture and its neural underpinnings. However, in natural settings reality confronts organisms with problems that are not neatly compartmentalized. Thus, a critical problem in memory research that still needs to be addressed is how distinct types of memories are ultimately integrated. Here we demonstrate how two memory networks, the hippocampus and dorso-lateral striatum, may accomplish such a goal. The hippocampus supports memory for facts and events, collectively known as declarative memory and often studied as spatial memory in rodents. The dorso-lateral striatum (putamen in primates) provides the basis for habits which are assessed in stimulus-response types of tasks. Expanding previous findings, the current work revealed that the hippocampus and dorso-lateral striatum use time and space in distinct and largely complementary ways to link spatial and habitual representations. Specifically, the hippocampus supported both types of memories when they were formed in temporal juxtaposition even if the learning took place in different environments. In contrast, the lateral striatum supported both types of memories if they were formed in the same environment even if at temporally distinct points. These results reveal for the first time that by using fundamental aspects of experience in specific ways, the hippocampus and dorso-lateral striatum can transcend their attributed roles in information storage.

**SIGNIFICANCE STATEMENT:** The current paradigm in memory research postulates that different types of memories reflected in separate types of behavioural strategies result from activity in distinct neural circuits. However, recent data have shown that when rats concurrently acquired in the same environment hippocampal-dependent spatial navigation and striatal-dependent approach of a visual cue, each of the two types of memories became dependent on both the hippocampus and dorso-lateral striatum. The current work reveals that the hippocampus and dorso-lateral striatum utilize distinct and complementary principles to integrate different types of memories in time and space: the hippocampus links memories formed in temporal proximity, while the lateral striatum links memories formed in the same space.

## INTRODUCTION

The multiple memory systems theory postulates that different types of memories result from the activity of distinct brain circuits with different properties and dynamics. The operational principle of the memory systems is considered to be independent parallelism, meaning that although information flows through all memory networks at the same time, processing within a given system occurs autonomously and each network supports one kind of memory (White, Packard, & McDonald, 2013; Squire, Knowlton, & Musen, 1993; Squire, 2004). This principle was formulated after numerous experiments in both animals and humans had shown that damage to one circumscribed memory circuit caused deficits in one of two behavioural tasks, each testing a distinct type of memory, while damage to a different memory circuit resulted in the opposite pattern. Based on numerous experiments of this kind, it is widely accepted that declarative memory, which includes spatial representations, is selectively dependent on a neural network centred on the hippocampus (HPC; Scoville & Milner, 1957; White & McDonald, 2002; Packard, Hirsh, & White, 1989; Squire & Zola-Morgan, 1991; McDonald & White, 1993; Eichenbaum, 2000; Aggleton & Pearce, 2001), while habits, which include stimulus-response (S-R) behaviours, require an intact dorso-lateral striatum (DSL; Squire, 2004; Knowlton, Mangels, & Squire, 1996; Packard & McGaugh, 1996; McDonald & White, 1994; Devan & White, 1999; Yin & Knowlton, 2004).

Recent data, however, have shown that the HPC and DSL can each support memories generally thought to be independent of these structures (Jacobson, Gruenbaum, & Markus, 2012; O’Reilly, Alarcon, & Ferbinteanu, 2014; Ferbinteanu, 2016). In the most recent of these studies(Ferbinteanu, 2016), when rats concurrently learned a spatial navigation and a cue response task on the plus maze in the same context (understood here as the environment within which the animal behaves), lesions of the HPC and DSL no longer resulted in the expected dissociation effects, which were still demonstrated if different groups of animals learned either of the two tasks alone. Instead, HPC and DSL lesions each impaired both spatial and response memory. Lesions of the medial striatum (DSM), a structure thought to be involved in flexible behaviour or behaviour based on action outcome associations (Ragozzino, Mizumori, & Kesner, 2002; Lee, Andre, & Pittenger, 2014; Yin, Ostlund, Knowlton, & Balleine, 2005), also impaired performance in both tasks, but in this case, the impairment occurred regardless of training parameters. These findings indicated that when spatial and response learning occur concurrently and in the same context, the HPC and DSL can be involved in behaviour that is incongruous with their otherwise known functions. Thus, these structures can engage in functional coupling to integrate memories of different kinds. If HPC-DSL functional coupling is based on a nonspecific, general process that occurs in memory networks when distinct types of learning occur in temporal proximity or constant environment, then separating the learning experiences in either time or space would presumably reverse the coupling, and the behavioural contributions of the two memory structures would shift towards the typical dissociation present when the animals learn only one task at a time. To test this hypothesis, the current experiment evaluated the memory deficits caused by HPC, DSL and DSM lesions when spatial and response learning were separated either in space but not time, or in time but not space (Fig. 1). One group of rats was trained concurrently in spatial navigation and cue response tasks, but each task was learned in a dedicated context (a condition referred to below as 2Contexts); this procedure separated the two kinds of learning in space but not time. A second group of rats learned the same tasks in one environment, but training on each task occurred during distinct days (a condition referred to below as 2Days); this procedure separated the two kinds of learning in time but not space. After reaching a set performance criterion, animals received selective HPC, DSL, DSM or sham lesions and were subsequently tested for retention following the same procedure as during training. Because rats trained in S-R tasks should be able to detect changes in context (McDonald, King, & Hong, 2001), the contexts were swapped during the last day of testing in the 2Contexts condition.

**Figure 1.**
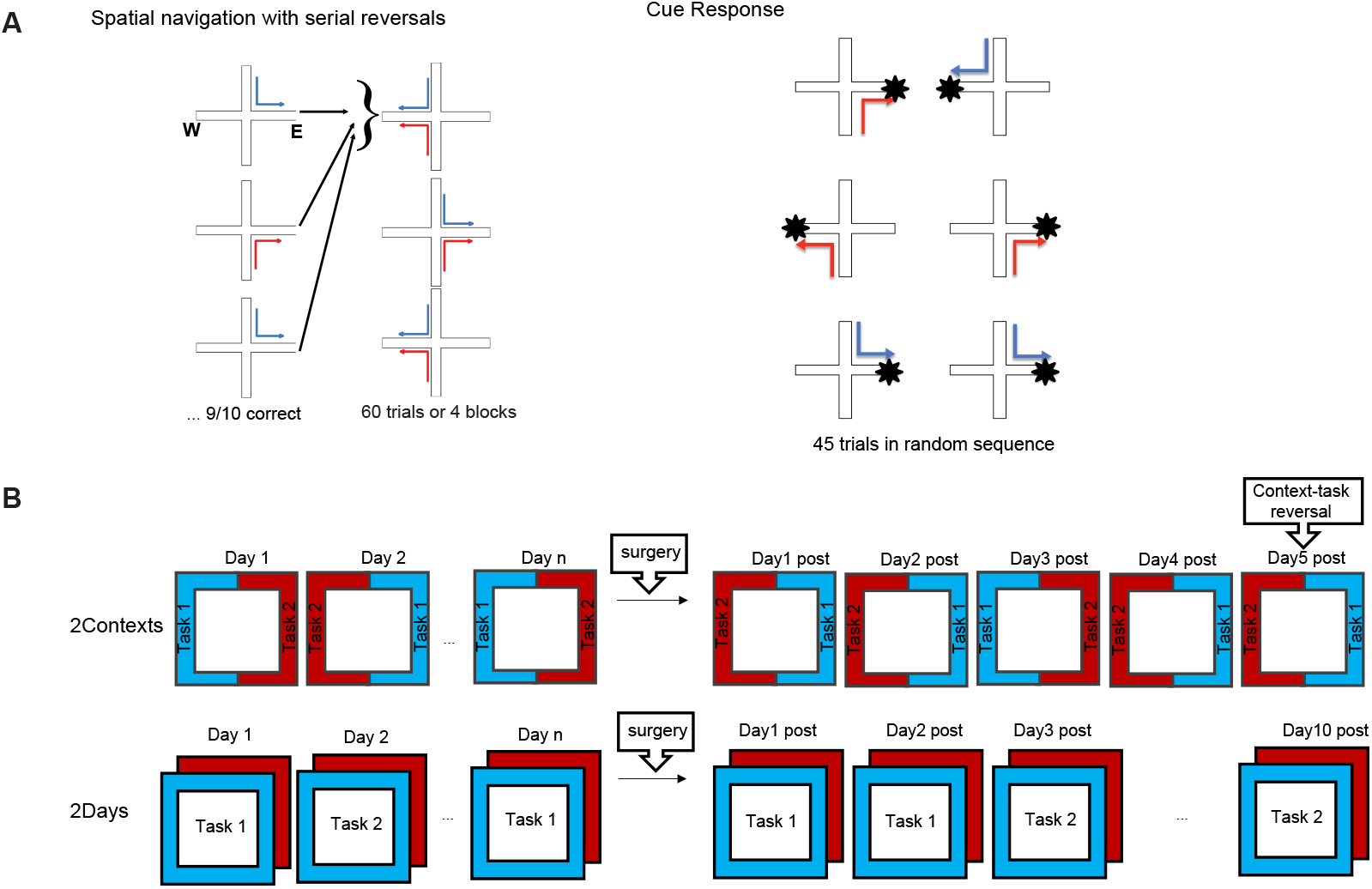
Behavioral paradigm and experimental design. A. The experiment used two distinct tasks, spatial navigation with serial reversals (left) and cue response (right). In both tasks the animals started either in the north or south arm of a plus maze apparatus, and hand to find food in either the east or the west arms. In the spatial navigation task, the food could be found based on its location, which was consistent in until the rat chose the correct arm 9 times correct in a sequence of 10 trials. At that point, the location of the food was switched to the other arm and a new block of trials started. Each session was constituted of 4 blocks to a total of maximum 60 trials. In the cue response task, the food was placed on a white flag whose position was randomly varied between the two goal arms to a total of 45 trials. The design ensures that the behavioral strategies suitable for solving each of the two tasks are mutually exclusive. B. Two training conditions, 2Contexts and 2Days, were used. In the 2Contexts condition, rats were trained/tested in both tasks during the same session, but each task was set in a dedicated context. On the 5th (last) day of retention testing, tasks and contexts were swapped (context-task reversal). In the 2Days condition, each rat was trained/tested in one context where it ran one task daily switching randomly between spatial and response learning across days.

## MATERIALS AND METHODS

### Subjects

Male Long–Evans rats (300-350g, 4-6 months old, Envigo) were individually housed (12-hour light cycle) and tested during the day. The animals were acclimatized to the colony, randomly assigned to one of four groups (DSL lesion, HPC lesion, DSM lesion, or sham) and food deprived to no more than 85– 90% of *ad libitum* body weight and kept at this standard throughout the experimental procedure. A total of 63 animals (32 rats for the 2Contexts condition and 31 rats for the 2Days condition) were included in this study, but only data from animals with lesions restricted to the intended areas were incorporated in the final analysis: DSM and DSL, 6 rats/group in either training condition; HPC 7 rats/group the 2Contexts condition and 8 rats/group in the 2Days condition. Sample size was determined based on previous work (Ferbinteanu, 2016) which showed that the same type of lesions as in the current experiment resulted in large differences relative to control groups. After training, animals within each training condition were randomly assigned to one of four groups: sham; HPC lesion; DSL lesion; and DSM lesion. All procedures were approved by the SUNY Downstate Medical Center Animal Care Committee (protocol 15-10452). The investigator was not blinded to group allocation, training-testing procedures, or during data analysis.

### Apparatus

The two plus mazes were made of grey polyvinyl chloride (PVC) and elevated 91 cm from the floor of two distinct rooms that each contained several visual cues and were illuminated distinctly. Each of the 4 arms was 61 cm long and 6.3 cm wide. A grey PVC block (30.4 cm high, 6.3 cm wide, 15.2 cm deep) was used to block the start arm that was not in use for that trial. In each case, a rectangular waiting platform (32cm × 42cm) was placed next to the maze. In the cued version of the task, a white visible flag also made of PVC was used to indicate the location of the food in the maze; during the spatial version of the task, the cue was placed on a table in the room and not in the maze.

### Behavioural Training and Testing

#### Experiment 1: Learning two types of tasks concurrently but in distinct contexts

One set of animals was trained in the two tasks on the same day, but each task was consistently associated with a dedicated environment. Each animal underwent one day of habituation, in which it was placed in each of the plus mazes in the presence of food; the visible flag was present in the maze in which the animal would then run the cue response task. Within each lesion group, the environments were counterbalanced across animals, and the tasks were learned in random order across days; once an animal completed a session in one environment, it was placed in the home cage and transferred to the second environment, where it underwent the second session. Training continued until the rats reached a criterion of 20% or less errors for two consecutive days in both tasks, after which they were assigned to one of four groups: HPC lesions, DSM lesions, DSL lesions, and sham controls. After a recovery interval of 5-10 days, retention was evaluated for 5 consecutive days. To test whether the animals associated a task with its dedicated environment, on the last day of post-lesion testing, each animal performed each of the two tasks in the ‘other’ environment.

#### Experiment 2: Learning two types of tasks in the same context but at distinct times

A second group of animals was trained in the same two tasks, but in this case, each animal was trained in only one context and learned the two tasks on different days. An example of a training session sequence is cue-cue-spatial-cue-spatial-cue-spatial-cue-cue-spatial-spatial-spatial-cue-spatial-cue-spatial-cue-cue. The training continued until the animal reached the 20% or less error criterion on two consecutive days for each of the two tasks, after which the animals were randomly assigned to the four lesion subgroups. To avoid a context-specific effect, the same two environments as in Experiment 1 were used in a counterbalanced manner (thus, in a lesion subgroup of six animals, three would be trained/tested in one environment and the remaining three in the other), as were the context/task of the first post-lesion retention test. After a recovery interval of 5-10 days, retention was evaluated for 10 days assigned randomly to 5 spatial and 5 response tests.

#### Spatial navigation and cue response tasks

All animals were pre-exposed to the maze in the presence of food for two consecutive days and then trained to walk from either the North or the South start arms to the end of West or East goal arms to obtain half a Fruit Loop. Between trials, the rats were placed on the side platform to wait for the next trial. Entry with all four paws into the unrewarded arm defined an *error*, which the rat was allowed to correct. Training procedures followed previously published protocols (Ferbinteanu, 2016) and utilized a spatial navigation task with serial reversals and a cue-response task (Fig. 1). In both cases, the start and the goal arm were selected based on a pseudorandom sequence of 60 trials with ? 3 consecutive repetitions of the same type of journey (NE, NW, SE or SW). *Spatial task.* In the spatial task, the animal was rewarded for remembering spatial location. The position of the food was kept constant until the rat entered the correct goal arm in 9 of 10 consecutive trials. At that point, the other goal arm was baited and a new block of trials began. If the animal did not reach the criterion in a maximum of 15 trials, the location of the food was changed automatically, to avoid unbalanced reinforcement of any specific goal arm. Alternating trial blocks continued up to either 4 blocks or 60 total trials. *Cue task.* In this case, the rats had to remember an association between the visible cue, whose location was rendered irrelevant by changing the start and goal positions based on a pseudorandom sequence, and a motor response, which was the walk towards the cue. Animals received 45 trials in each session, a number approximately equal to the number of trials necessary to complete four blocks of trials in the spatial task. Thus, each goal arm was approximately equally rewarded both within and across tasks.

#### Lesions

At the end of the training phase, animals were randomly assigned to one of four groups: sham controls, HPC lesion group, DSL lesion group, and DSM lesion group. Rats were anaesthetized with isoflurane and diazepam (10mg/kg). Atropine (5 mg/kg body weight) was also administered to avoid fluid accumulation in the respiratory tract. Neurotoxic lesions were made by injecting either a solution of 5mg/ml NMDA in phosphate buffer (Tocris; pH 7.4) or a solution of quinolinic acid (Tocris; 25mg/ml in phosphate buffer titrated with sodium hydroxide to pH 7.4) through a 30-gauge cannula attached to a minipump (0.2 μl/min; New Era Pump Systems, Inc., Model NE-4000). At the end of each injection, the cannula was left in place for 3 mins, retracted 0.5 mm and left in this location for 2 mins, after which it was slowly and completely retracted. The coordinates of each injection and the volumes injected are presented in Table 1. To prevent seizure development, a second, i.p., injection of diazepam (10 mg/kg body weight) was administered prior to neurotoxin infusion, and the animals were monitored until completely awake and active in their home cages. Sham animals were anaesthetized, incised and sutured.

**Table 1.**
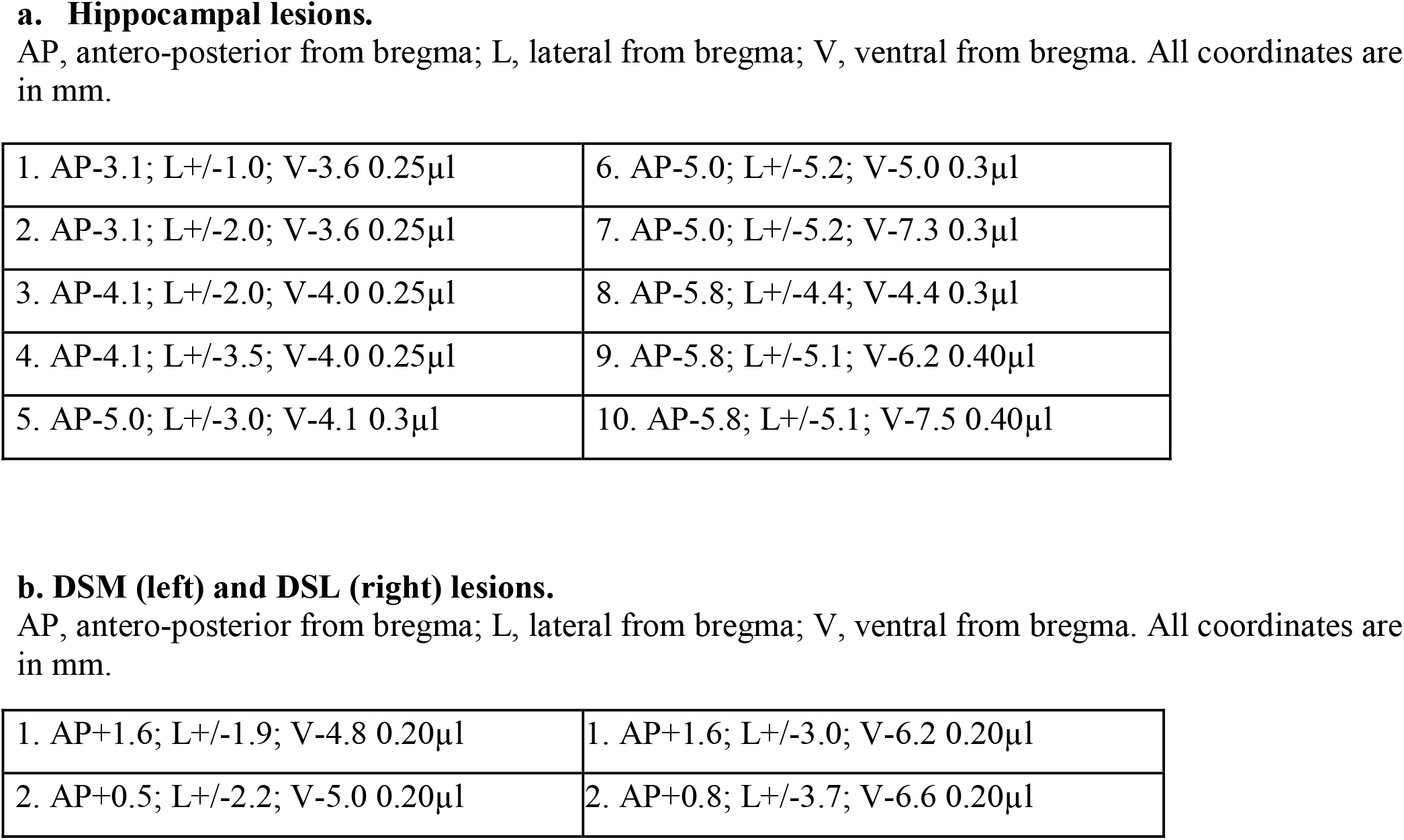
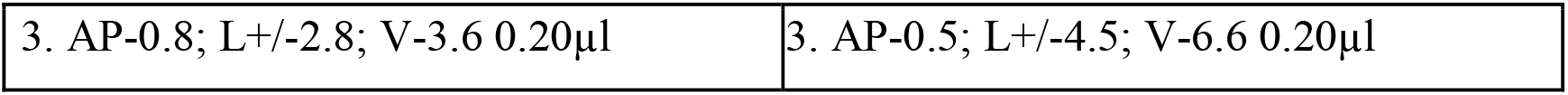
Lesion coordinates.

|  |  |
| --- | --- |
| 1. AP-3.1; L+/-1.0; V-3.6 0.25μl | 6. AP-5.0; L+/-5.2; V-5.0 0.3μl |
| 2. AP-3.1; L+/-2.0; V-3.6 0.25μl | 7. AP-5.0; L+/-5.2; V-7.3 0.3μl |
| 3. AP-4.1; L+/-2.0; V-4.0 0.25μl | 8. AP-5.8; L+/-4.4; V-4.4 0.3μl |
| 4. AP-4.1; L+/-3.5; V-4.0 0.25μl | 9. AP-5.8; L+/-5.1; V-6.2 0.40μl |
| 5. AP-5.0; L+/-3.0; V-4.1 0.3μl | 10. AP-5.8; L+/-5.1; V-7.5 0.40μl |
| 1. AP+1.6; L+/-1.9; V-4.8 0.20μl | 1. AP+1.6; L+/-3.0; V-6.2 0.20μl |
| 2. AP+0.5; L+/-2.2; V-5.0 0.20μl | 2. AP+0.8; L+/-3.7; V-6.6 0.20μl |
| 3. AP-0.8; L+/-2.8; V-3.6 0.20µl | 3. AP-0.5; L+/-4.5; V-6.6 0.20µl |

#### Behavioural Data Analysis

Percent performance error was calculated for each rat during each day of testing, and a mean was calculated for each group during each day. All analyses were performed using SAS version 9.2 (SAS Institute, Inc., NY). Differences in performance were assessed using two-way mixed model analyses. Time (in days) and lesion group were entered into the model as categorical independent variables, and performance as percent error on repeated time points was entered as a dependent variable. The degrees of freedom were computed according to the Satterthwaite formula, which takes into consideration the variance within the group along with the sample size and is robust against heterogeneity of variance. Where overall analyses indicated significant differences, differences in performance between each lesion group and the sham group were further investigated. The effects of the reversal test (5^th^ post-lesion day in the 2Contexts condition) were assessed by using a two-tailed t-test to evaluate for each group the behavioural performance during the test relative to the performance during the previous day.

#### Tissue Preparation and Processing for Histology

Rats were overdosed with isoflurane (administered in a closed environment) and perfused transcardially with normal saline followed by 10% formalin for tissue fixation. Coronal sections (40 μm) were cut on a cryostat and stained with cresyl violet to evaluate the extent of the lesion.

#### Lesion Assessment

Brains were coronally sectioned. Each 4^th^ section in the striatum lesion groups and each 5^th^ section in the hippocampal lesion groups was mounted on a microscope slide to be used for lesion evaluation. We first visually inspected the sections under the microscope and traced each lesion on a set of histological plates (Paxinos & Watson, 1988). Data from animals whose lesions were not sufficiently inclusive (i.e., not encompassing most of the targeted area) and selective (i.e., extending bilaterally to significant portions of other brain areas) were excluded from further analysis. The lesions that met the criteria underwent quantification analysis. All slides incorporated in the analysis were scanned using a slide scanner (Aperio AT2, Leica Biosystems) to generate digital images of the sections. Dedicated software (ImageScope, Leica Biosystems) was then used to visualize and measure for each section the area of preserved hippocampal or striatal tissue. This procedure was also performed for brain tissue from four control animals for the dorsal striatum and four control animals for the hippocampus. Ten sections were selected from each animal in the DSM and DSL groups and 14 sections in the HPC groups (approximately half the total number of sections) so that approximately equivalent levels on the antero-posterior axis were captured in the analysis across animals. The values of the preserved tissue areas were then summed to estimate the total healthy tissue for each animal. (Because the brain tissue was sampled at regular intervals of 200 μm for all HPC tissue and 160 μm for all striatal tissue in control and lesion animals alike, multiplying by the constant distance between sections was not necessary to obtain an estimate of the lesion size as a % of the total structure.) The totals for the control animals were averaged for the dorsal striatum and hippocampus, and the resulting values were considered as 100% in the computation that evaluated the lesion size. For each lesioned animal, %healthy tissue was computed by dividing the area of healthy tissue in their brain by the corresponding area in the control brain; the %lesion was then obtained as the difference between the %healthy tissue and 100%. For each lesion group, we compared the % damage across the three experimental conditions using an unpaired 2 tailed t-test (The SAS Institute).

## RESULTS

### Lesions encompassed the intended brain areas

For each animal, the lesion’s selectivity was evaluated; only data from animals with damage restricted to the intended brain areas were considered for further analysis. Neurotoxic cortical damage was small and typically only the mechanical damage at the canula insertion point was present. All tissue with signs of gliosis was considered lesioned, regardless of whether the principal neurons might have been spared at the periphery of the affected area (Fig. 2A, orange arrowheads). For the HPC lesion groups, all areas of HPC proper (dentate gyrus, CA3, CA2 and CA1) incurred large, almost complete, damage in both dorsal and ventral portions of the two structure (Fig. 2, top rows). In general, the ventral tip of the HPC was spared, as well as the ventral blade of the dentate gyrus at the most anterior HPC pole. There was minor damage to the subiculum in the anterior areas but no damage was found in the entorhinal cortex or other nearby cortical regions. Two animals in each of the two HPC lesion groups had small cortical damage. There was small unilateral damage in the thalamus in one rat in the 2Days condition resulting from a deep penetration of the canula, which was not accompanied by neurotoxic damage; the data of this animal did not show marked differences from the data of the rest of the subjects in the same group. In the dorsal striatum of the rat, the caudate and putamen, which correspond to medial and lateral areas, are fused. Because the effects of small lesions preclude clear functional interpretations if animals recover from memory deficits during retention tests, here we aimed for lesions as inclusive as possible, that would conform to known topography of anatomical connections (McGeorge & Faull, 1989). For DSL lesion groups, 3 animals in the 2Contexts condition and 4 animals in the 2Days condition had small unilateral damage to claustrum, endopiriform cortex, lateral orbital cortex, and globus pallidus. For DSM lesion groups, 1 animal in the 2Contexts condition and 2 animals in the 2Days condition had small damage to globus pallidus. Overall, the lesions distinguished well between the medial and lateral areas of the dorsal striatum with overlap restricted to the most anterior and dorsal regions of the striatum (Fig. 2B). Lesion size was not significantly different across training conditions in any of the lesion groups (DSM: t_10_ = 1.678, p = 0.1241; DSL: t_10_ = 0.166, p = 0.8711; HPC: t_13_ = 1.073, p = 0.3084; Fig. 2A right inset, Fig. 2B).

**Fig. 2A.**
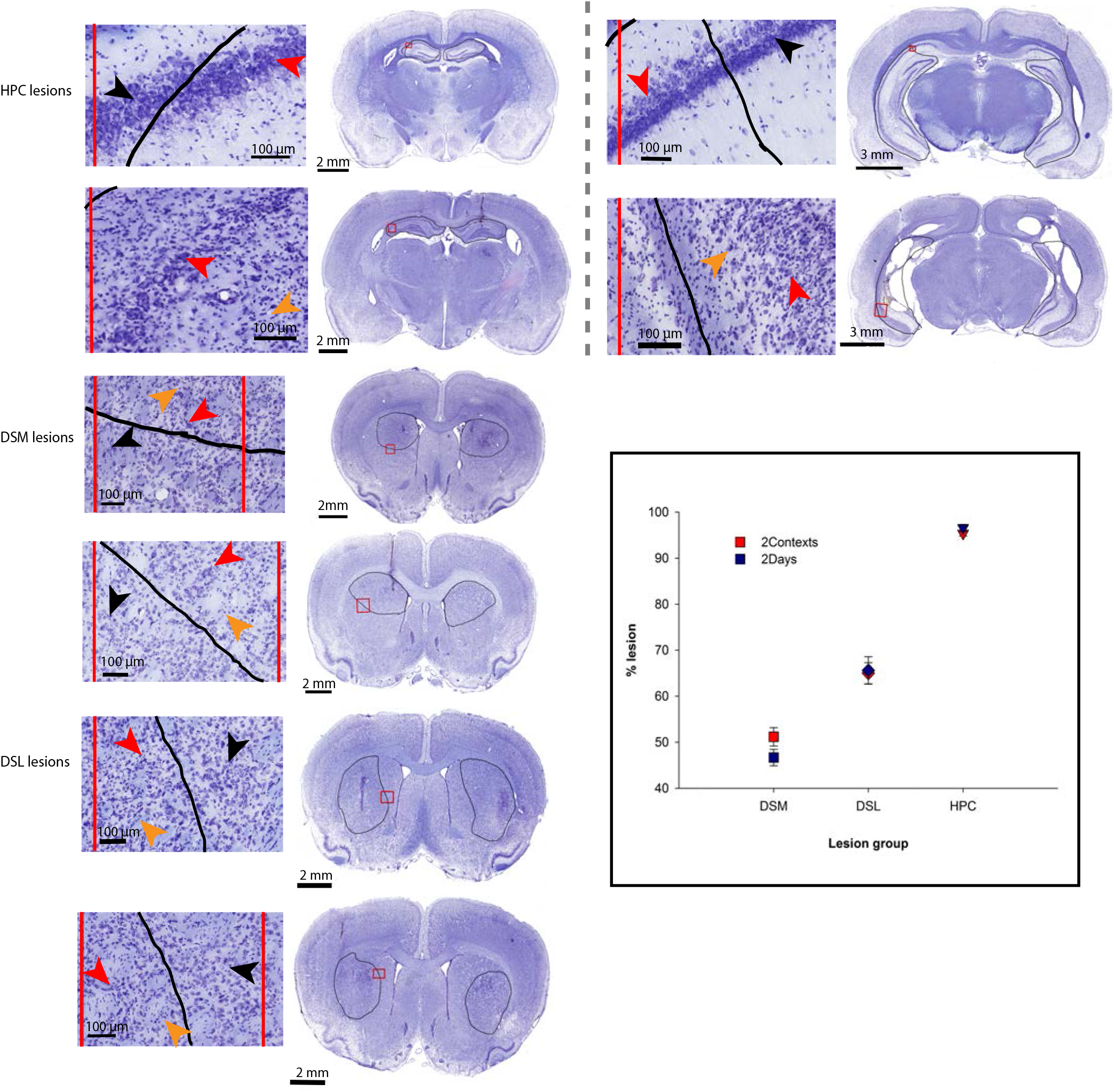
Lesions were localized in the targeted areas. Each of the six lesion groups (HPC, n = 7 and n = 8, respectively; DSM, n = 6 and n = 6, respectively; and DSL n = 6 and n = 6, respectively groups in 2Contexts and 2Days training conditions) is represented by one lesion illustration. Top two rows (2 different rats) show sections in dorsal (left) and ventral (right) HPC from the same animal. Photomicrographs of coronal sections through the brain at low magnification indicate the lesion area (circled in black). Enlarged parts of the small area within the red square are presented to the left in each case to enable comparison between degenerating or dead cells (red arrowheads) and healthy cells (black arrowheads). For both DSM and DSL, tissue with any signs of gliosis (such as astrocytic invasion among apparently normal cells, orange arrowheads) was marked as lesioned. Similarly, HPC lesions included areas where neurons were either missing or degenerating, or astrocytes were seen in large numbers. The inset presents the results of the quantification analysis. Lesion size was not significantly different across training conditions in either of the lesion groups (DSM: t_10_ = 1.678, p = 0.1241; DSL: t_10_ = 0.166, p = 0.8711; HPC: t_13_ = 1.073, p = 0.3084).

**Fig. 2B.**
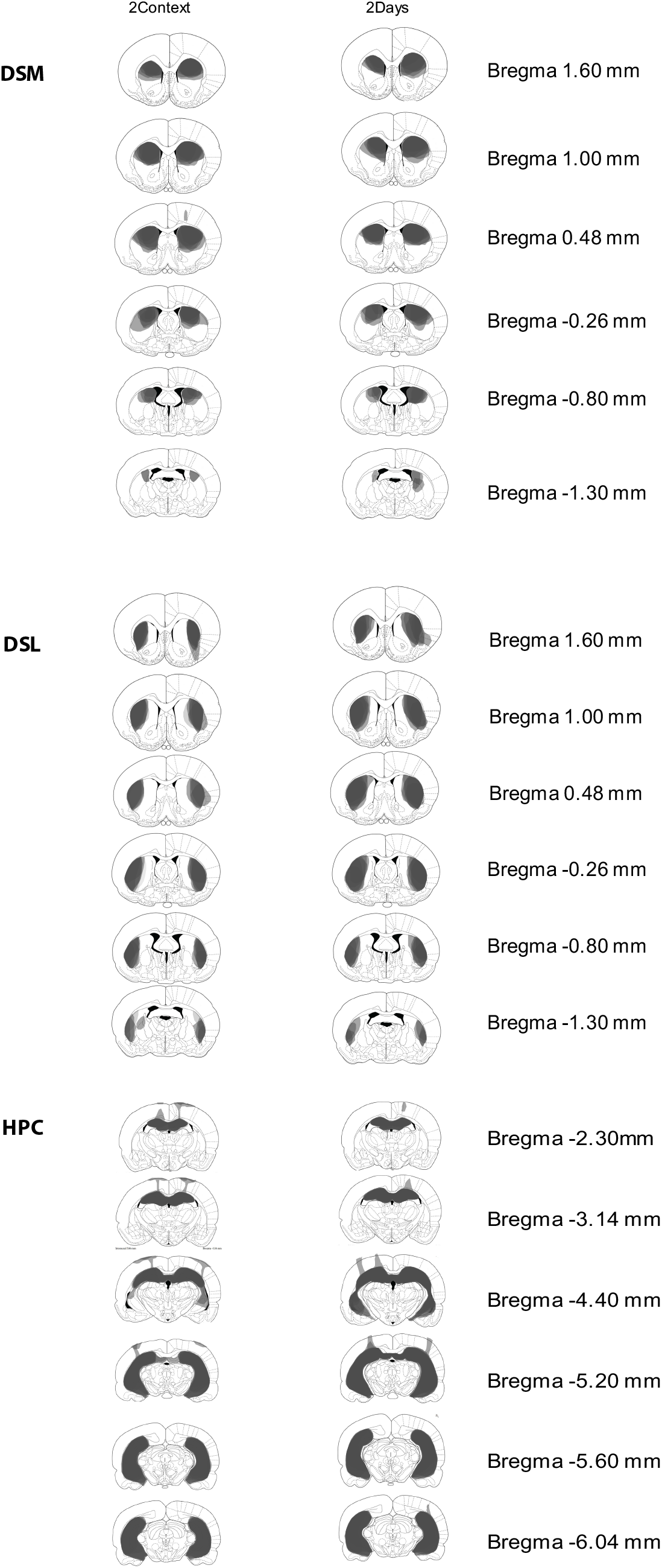
Extent of the lesions. For all animals whose data were included in the analysis, lesioned areas were marked and superimposed on the same set of corresponding diagrams. Areas lesioned in all animals of a group show as the darkest gray, areas lesioned in only one animal are shown as lightest gray. The brain tissue was damaged at the insertion point of the injection needle in all cases; the figure shows only damage larger than what would be expected from the 30 Ga canula. All tissue with signs of gliosis was considered as part of the damaged area.

### DSL supported spatial memories acquired in the same environment as response memories; HPC supported response memories acquired concurrently with spatial memories

Rats with DSL lesions showed a significant spatial memory deficit relative to controls throughout the 5 days of retention testing only when spatial and response memories were acquired in the same environment (Fig. 3A, left; Table 2a). Directly comparing the spatial performance of animals with this type of lesions in the two training conditions highlighted the large difference that training procedure caused on performance (F_(1, 47.9)_ = 84.14, p < .00001; Fig. 3B, upper left panel). At the same time, as expected, DSL lesions impaired response memory regardless of training procedure (Fig. 3A, right; Table 2b; for more on the recovery from the memory deficit in the 2Contexts condition see below). Thus, while DSL supports a rigid habitual response to a single cue in general, it can also become involved in a behaviour illustrative of declarative memory if spatial and response learning occur in the same space.

**Figure 3.**
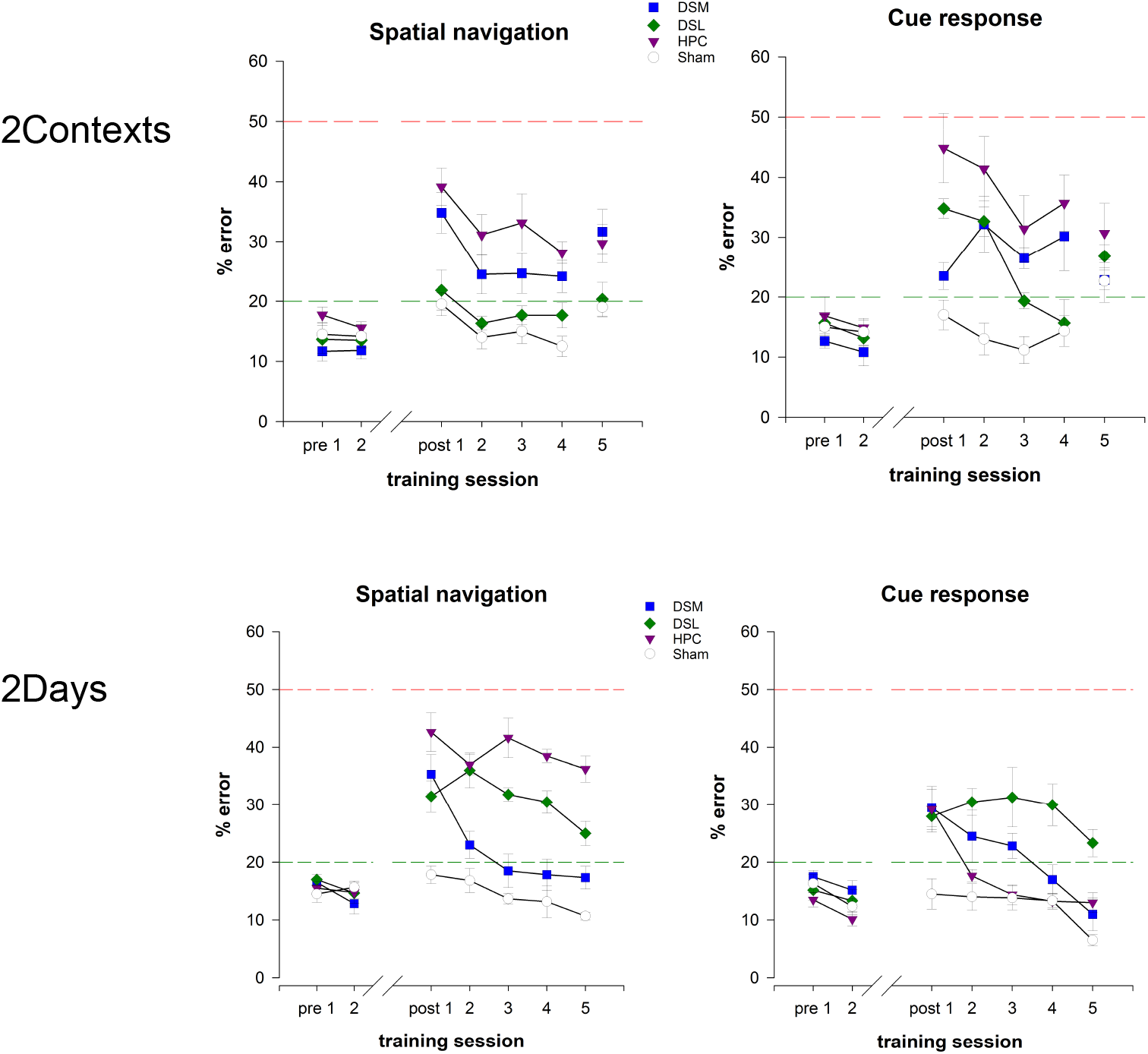
Behavioral performance for all groups in the two training conditions. **A.** Data organized to facilitate comparisons among lesion groups within the same training condition and performance during the reversal test in the 2Context condition. All rats performed well before the surgery and sham animals (n=6/group) continued to do so after the surgery. In contrast, lesioned animals exhibited memory deficits. In the 2Contexts condition, the DSL group (n = 6) had normal spatial and impaired response memories while the HPC group (n=7) made many errors in both spatial and response tasks. In the 2Days condition, the pattern reversed: the DSL group (n=6) had both spatial and response memory deficits, while the HPC group (n = 8) showed impaired spatial but largely normal response memory (see 2B. and text for details of performance during first day post lesion). The DSM groups (n = 6/group) showed various degrees of memory deficits throughout retention testing. Context reversal disrupted the performance of sham animals. Rats with HPC lesions did not alter their performance, the DSL lesion group performed worse in the response task, and the DSM lesion group performed worse in the spatial task. For all graphs, vertical axes show percentage error in behavioral performance (mean +/− SEM) and horizontal axes show training/testing sessions, with the break marking the surgery point. The green horizontal lines at 20% indicate the criterion threshold. The red horizontal lines at 50% indicate chance performance level.

**Figure 3B.**
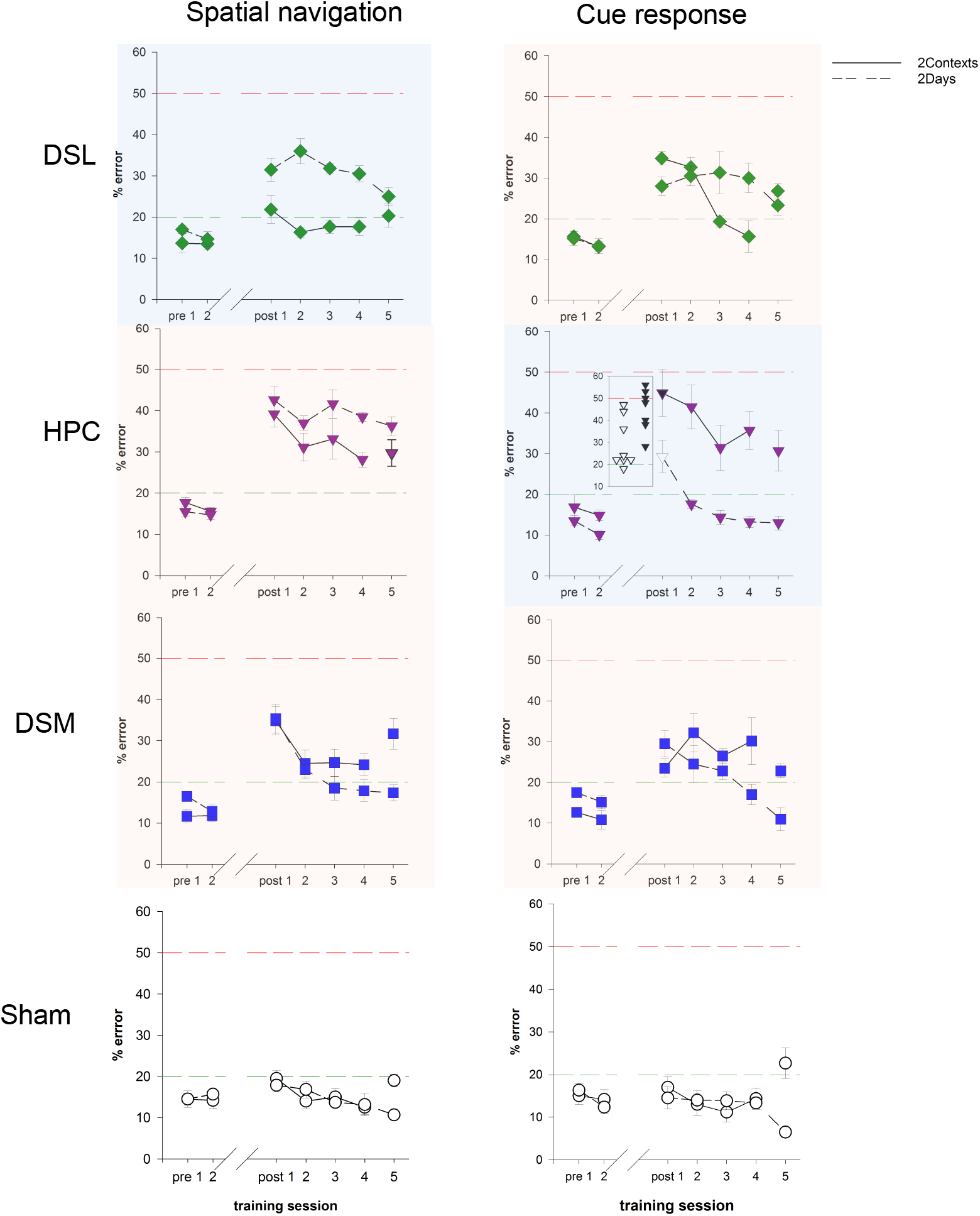
Same data as in A organized to reveal effects of training conditions. DSL was critically involved in response memory overall, but also supported spatial navigation when spatial and response learning occurred in the same context during different days (upper row, right). HPC was critical for spatial navigation overall, but additionally supported cue response when spatial and response learning occurred concurrently in different contexts (second row, left). The inset shows individual data points during the first day retention testing for the 2Contexts (black) and 2Days (white) conditions. Five of the 8 animals in the 2Days condition were close to criterion level. In contrast, 6 out of 7 animals in the 2Contexts condition showed severe impairment. Sham animals performed well regardless of training condition. In all cases, statistical analyses included only data from days 1-4 post-lesion training (i.e., excluded the context/task reversal test). Blue highlights indicate the data that do not conform to the current thinking on memory organization, orange highlights indicate the data that do.

**Figure 3C.**
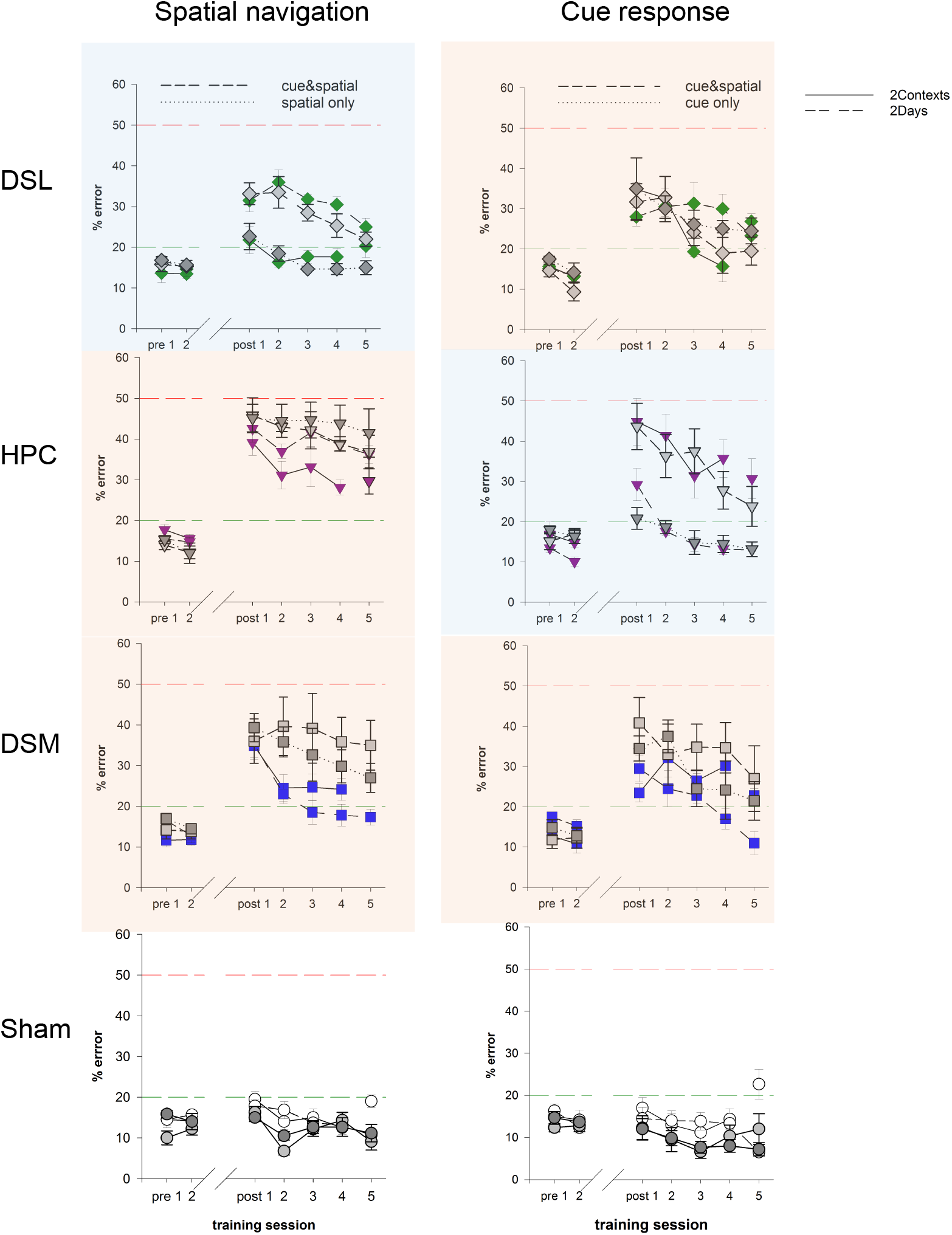
Same data as in A and B (in color), combined with previous data (Ferbinteanu, 2016) illustrating the effects of DSL, HPC and DSM lesions on spatial and response memories after training in one task only (spatial only and cue only; dark gray), or after training in the two tasks concurrently in the same context (cue&spatial; light gray). The effects of training paradigm on the performances of DSL groups in spatial navigation and HPC groups in cue response are similar in the two experiments (blue highlights). Thus, the time factor can account by itself for the effect on response memory caused by HPC lesions when both time and space factors were involved. Similarly, the space factor can account by itself for the effect on spatial memory caused by DSL lesions when both time and space factors were involved. In contrast, the performances of DSL lesion groups in the cue response task and HPC lesion groups in the spatial navigation task varied between this and previous experiments, as did the performance of the DSM lesion groups overall (orange highlights). The behavior of sham animals remained consistent across training protocols.

**Table 2.**
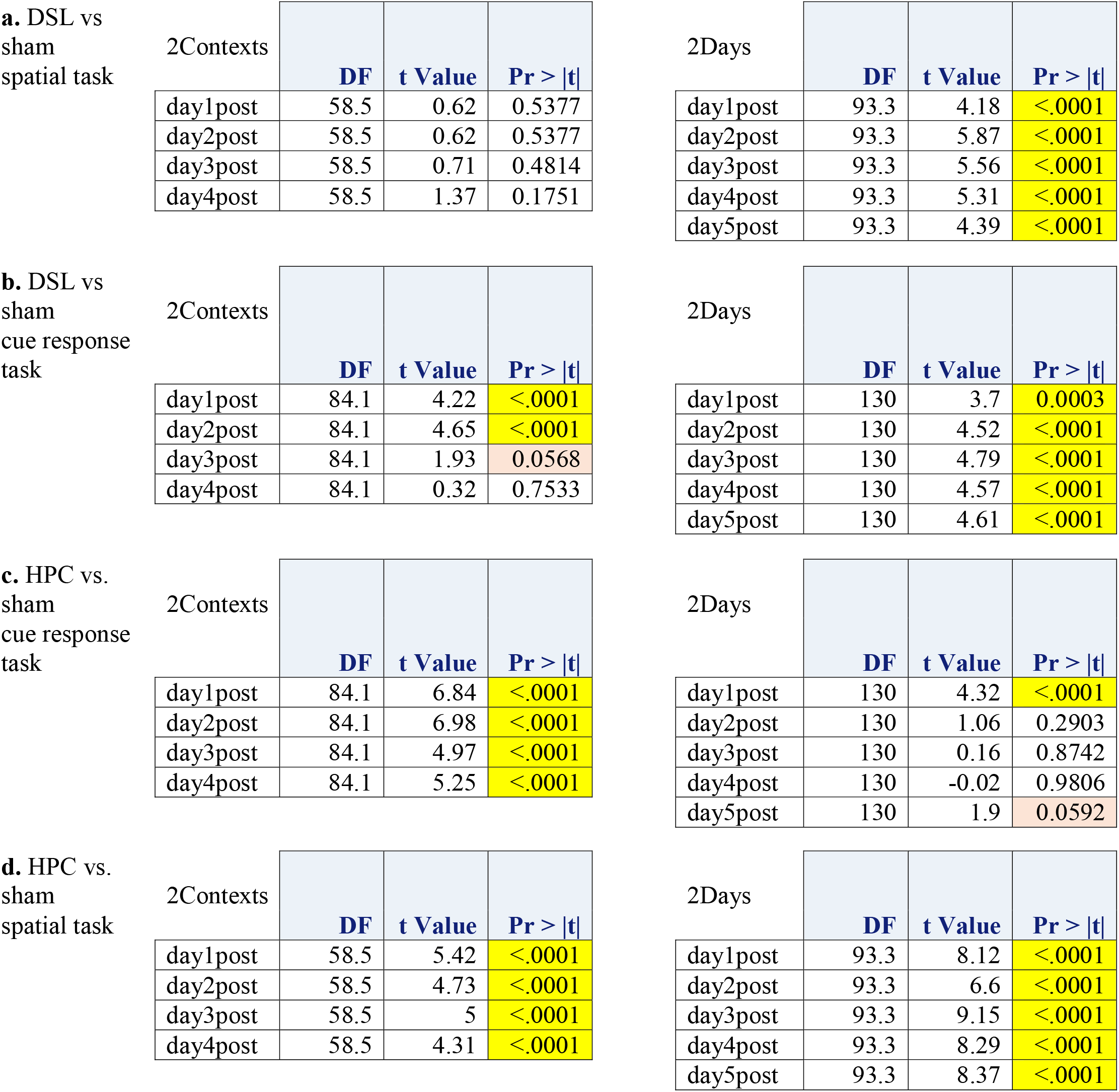
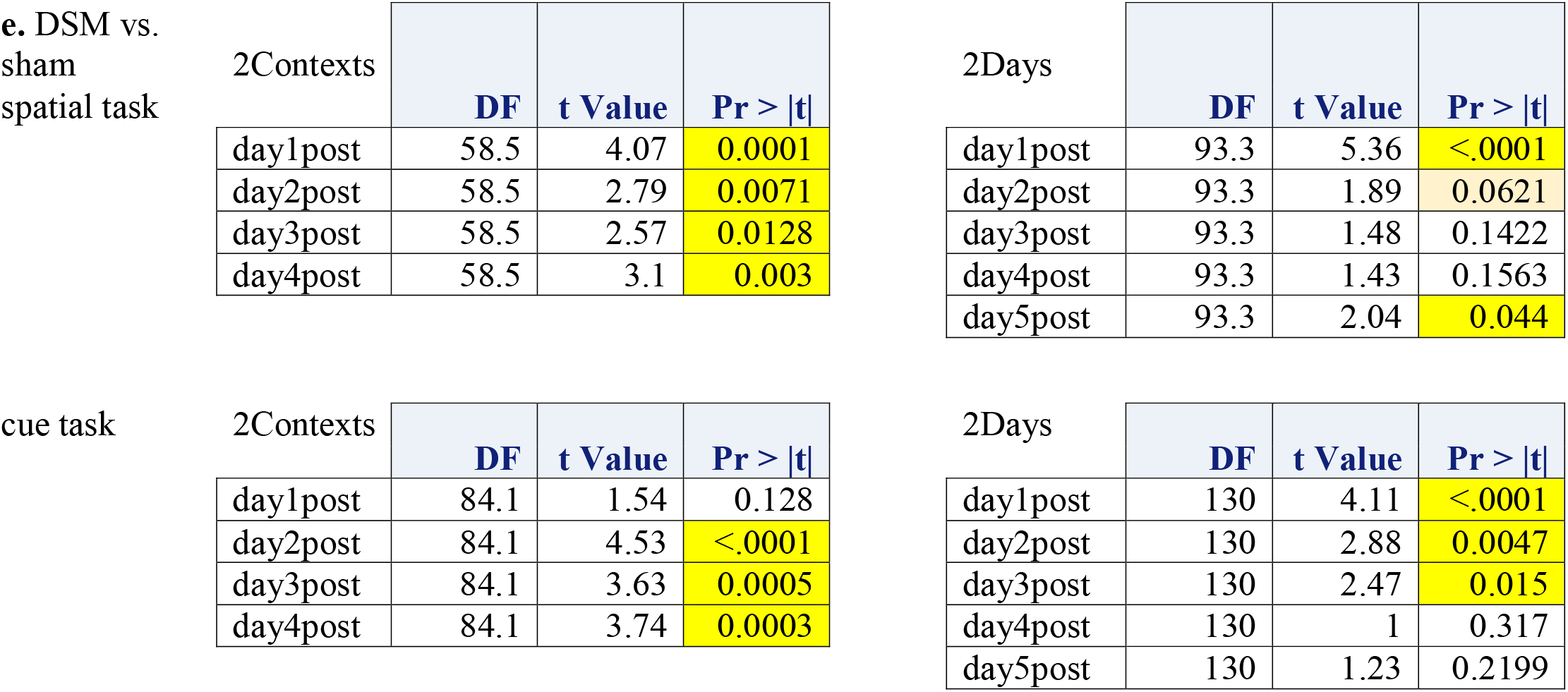
Response and spatial memory deficits in DSL and HPC lesion groups, respectively. Each section list the results of t-tests comparing performance of animals with DSL, HPC, or DSM lesions to the performance of corresponding sham groups. The tests were a priori planned and run after the results of the two-way mixed analyses indicated highly significant main effects and interactions; the degrees of freedom were computed based on Welch-Satterthwaite equation. DF = degrees of freedom; yellow highlights = statistically significant; pink highlights = marginally significant

**a. DSL vs sham spatial task**
| 2Contexts | DF | t Value | Pr > \|t\| |
| --- | --- | --- | --- |
| day1post | 58.5 | 0.62 | 0.5377 |
| day2post | 58.5 | 0.62 | 0.5377 |
| day3post | 58.5 | 0.71 | 0.4814 |
| day4post | 58.5 | 1.37 | 0.1751 |

| 2Days | DF | t Value | Pr > \|t\| |
| --- | --- | --- | --- |
| day1post | 93.3 | 4.18 | <.0001 |
| day2post | 93.3 | 5.87 | <.0001 |
| day3post | 93.3 | 5.56 | <.0001 |
| day4post | 93.3 | 5.31 | <.0001 |
| day5post | 93.3 | 4.39 | <.0001 |

**b. DSL vs sham cue response task**
| 2Contexts | DF | t Value | Pr > \|t\| |
| --- | --- | --- | --- |
| day1post | 84.1 | 4.22 | <.0001 |
| day2post | 84.1 | 4.65 | <.0001 |
| day3post | 84.1 | 1.93 | 0.0568 |
| day4post | 84.1 | 0.32 | 0.7533 |

| 2Days | DF | t Value | Pr > \|t\| |
| --- | --- | --- | --- |
| day1post | 130 | 3.7 | 0.0003 |
| day2post | 130 | 4.52 | <.0001 |
| day3post | 130 | 4.79 | <.0001 |
| day4post | 130 | 4.57 | <.0001 |
| day5post | 130 | 4.61 | <.0001 |

**c. HPC vs. sham cue response task**
| 2Contexts | DF | t Value | Pr > \|t\| |
| --- | --- | --- | --- |
| day1post | 84.1 | 6.84 | <.0001 |
| day2post | 84.1 | 6.98 | <.0001 |
| day3post | 84.1 | 4.97 | <.0001 |
| day4post | 84.1 | 5.25 | <.0001 |

| 2Days | DF | t Value | Pr > \|t\| |
| --- | --- | --- | --- |
| day1post | 130 | 4.32 | <.0001 |
| day2post | 130 | 1.06 | 0.2903 |
| day3post | 130 | 0.16 | 0.8742 |
| day4post | 130 | -0.02 | 0.9806 |
| day5post | 130 | 1.9 | 0.0592 |

**d. HPC vs. sham spatial task**
| 2Contexts | DF | t Value | Pr > \|t\| |
| --- | --- | --- | --- |
| day1post | 58.5 | 5.42 | <.0001 |
| day2post | 58.5 | 4.73 | <.0001 |
| day3post | 58.5 | 5 | <.0001 |
| day4post | 58.5 | 4.31 | <.0001 |

| 2Days | DF | t Value | Pr > \|t\| |
| --- | --- | --- | --- |
| day1post | 93.3 | 8.12 | <.0001 |
| day2post | 93.3 | 6.6 | <.0001 |
| day3post | 93.3 | 9.15 | <.0001 |
| day4post | 93.3 | 8.29 | <.0001 |
| day5post | 93.3 | 8.37 | <.0001 |
| day1post | 58.5 | 4.07 | 0.0001 |
| day2post | 58.5 | 2.79 | 0.0071 |
| day3post | 58.5 | 2.57 | 0.0128 |
| day4post | 58.5 | 3.1 | 0.003 |

2Days
| 2Days | DF | t Value | Pr > \|t\| |
| --- | --- | --- | --- |
| day1post | 93.3 | 5.36 | <.0001 |
| day2post | 93.3 | 1.89 | 0.0621 |
| day3post | 93.3 | 1.48 | 0.1422 |
| day4post | 93.3 | 1.43 | 0.1563 |
| day5post | 93.3 | 2.04 | 0.044 |

cue task
| 2Contexts | DF | t Value | Pr > \|t\| |
| --- | --- | --- | --- |
| day1post | 84.1 | 1.54 | 0.128 |
| day2post | 84.1 | 4.53 | <.0001 |
| day3post | 84.1 | 3.63 | 0.0005 |
| day4post | 84.1 | 3.74 | 0.0003 |

2Days
| 2Days | DF | t Value | Pr > \|t\| |
| --- | --- | --- | --- |
| day1post | 130 | 4.11 | <.0001 |
| day2post | 130 | 2.88 | 0.0047 |
| day3post | 130 | 2.47 | 0.015 |
| day4post | 130 | 1 | 0.317 |
| day5post | 130 | 1.23 | 0.2199 |

Current data do not clarify whether the DSL formed its own context representation, directly accessed the context representation formed by the HPC, or accessed the HPC context representation at the level of control over the motor output. The results of the reversal test in the 2Contexts condition suggested that in this experiment, context representations were selectively dependent on the HPC because the performance of this lesion group was not altered by the context swap in either of the two tasks (Fig. 3A top row; Table 3).

**Table 3.** Results of the reversal test. The tables list the results of a priori planned t-tests comparing for each lesion group the performance during the reversal test in the 2Contexts condition relative to the preceding day. DF=degrees of freedom; yellow highlights = statistically significant; pink highlights = marginally significant

| spatial task | lesion | DF | t Value | Pr > \|t\| |
| --- | --- | --- | --- | --- |
|  | DSL | 25 | -1 | 0.3259 |
|  | DSM | 25 | -2.82 | 0.0093 |
|  | HPC | 25 | -0.64 | 0.5293 |
|  | sham | 25 | -2.44 | 0.022 |

| cue task | lesion | DF | t Value | Pr > \|t\| |
| --- | --- | --- | --- | --- |
|  | DSL | 25 | -2.44 | 0.022 |
|  | DSM | 25 | 1.6 | 0.1212 |
|  | HPC | 25 | 1.18 | 0.2485 |
|  | sham | 25 | -1.82 | 0.0802 |

The performance of the HPC lesion groups on the cue response task showed a pattern of impairment largely complementary to the one described above for animals with DSL lesions on the spatial task (Fig. 3A right). Specifically, these groups were consistently impaired in response behaviour when the rats formed spatial and response memories concurrently but in distinct environments (Fig. 3B, second row right; Table 2c). In the other training condition, when spatial and response memories were acquired in the same environment across different days, the HPC lesion group had a deficit in response memory only on the first day of retention testing. The result was due to the poor performance of 3 out of 8 animals (Fig. 3B, inset of second row, right hand panel). These animals’ rapid recovery of the memory deficit on the subsequent day suggested that an already existing cue-response association regained control over the motor output. Directly comparing response performance in animals with HPC lesions that underwent different training procedures underscored the large differences in the behaviour of these two groups (F_(1, 60)_ = 114.55, p < 0.0001). In contrast with the selective effect on response performance, HPC lesions consistently impaired spatial navigation (Fig. 3B, second row left; Table 2d). Thus, the HPC is critical for spatial representations in general, but it can additionally support response memories if they are formed in close temporal proximity to spatial memories.

### DSM supported flexible behaviour

Although the DSL and DSM are both parts of the dorsal striatum, the DSM is involved not in habitual, but flexible, goal-directed behaviours (Ragozzino, Jih, & Tzavos, 2002; Ragozzino et al., 2002; Yin, Knowlton, & Balleine, 2006; Yin et al., 2005; Gremel & Costa, 2013). The current results were in agreement with this idea, as DSM lesions generally impaired performance regardless of training conditions in both the spatial and response tasks. The degree of impairment varied with training procedure but the size of the effect was considerably smaller than for the other two types of lesions and stronger in the response than in the spatial task (Fig. 3A and 3B, third row; spatial navigation: F_(1, 47.9)_ = 3.57, p = 0.0650; cue response: F_(1, 47.9)_ = 5.11, p = 0.0283; Table 2e). This pattern of results is not surprising given that in each task, the animals have to adapt the body turn at the centre of the maze to their start position relative to the goal. The data also suggested that the DSM lesions may cause slightly more impairment in the 2Contexts condition in general, also not surprising because in this condition, the animals have to switch from one task to another within the same test session. Thus, the DSM supported both spatial navigation and habitual response to a discrete cue, and the data overall indicate that its contribution is consistent with a role in behavioural flexibility.

### The magnitudes of spatial and response memory deficits were modulated by training procedure

A large body of empirical data links the DSL to S-R behaviour and the HPC to spatial navigation. The results of the current experiment confirmed this idea but also indicated that for both structures, the magnitude of the memory deficit was modulated by the training condition (HPC: F_(1, 59.9)_ = 9.86, p = 0.0026; DSL: F_(1, 47.3)_ = 4.54, p = 0.0383; Fig. 3B, top row right and second row left, orange highlights). Animals with HPC lesions had a permanent spatial memory deficit regardless of how the task was learned. Animals with DSL lesions showed permanent impairment in response memory in the 2Days condition. In the 2Contexts condition, this lesion group showed initially a strong impairment during the first two days of retention testing which then declined rapidly and was eliminated in the next two days. Thus, in this case and in this case only, the remaining neural circuits (which do not include remnants of the DSL network) can rescue the response behaviour. As already described above, training condition also modulated the performance of the DSM lesion groups but note that in this case, the 2Contexts condition was consistently associated with more impairment, which was not the case with DSL and HPC lesions. In contrast, the training condition did not affect the proficient performance of normal animals (Fig. 3B, bottom row). Collectively, these findings showed that the specialized representations each of the three brain structures form can support the corresponding behaviours as a matter of degree, depending on training condition. If so, then it is of interest to know how the results from the current experiment compare to previous work (Ferbinteanu, 2016), when animals with similar types of lesions learned the same two tasks concurrently and in the same environment (that is, in a way that combined the spatial and temporal factors). Thus, the past and current results were plotted on the same graphs (Fig. 3C). Comparison between these two data sets revealed two facts. First, for both the DSL and HPC, the pattern of contribution to the task incongruous with that structure’s style of information processing—DSL to spatial navigation and HPC to cue response—were remarkably similar (Fig. 3C, blue highlights). Because the DSL groups performed the same in spatial navigation regardless of whether the rats learned the two tasks concurrently (previous data) or on distinct days (current data), it seems that the temporal factor did not modulate the ability of the DSL to link distinct memories based on space. Similarly, because the HPC groups performed the same on cue response regardless of whether the rats learned the two tasks in the same (previous data) or distinct (current data) contexts, it seems that the spatial factor did not impact the ability of the HPC to link distinct memories based on time (with the caveat mentioned above). This finding further supports the idea that the HPC and DSL integrate memories based not only on distinct, but also largely complementary functional principles. Second, the comparison between past and present results further confirmed that in tasks congruous with the type of representation each brain area forms, the lesion effects were modulated by the training protocol (Fig. 3C, orange highlights). The effects do not suggest a hierarchical organization such that the highest levels of deficit would result when spatial and temporal factors are combined (animals learn cue and spatial tasks together in the same context; previous experiment), next down would be the effects corresponding to one of the two factors (temporal or spatial by themselves; the present experiment) and least deficit would result when neither factor is present (animal learn only one task at a time). Thus, depending on learning circumstances, the same normal learned behaviour, currently thought of as being controlled by a set neurobiological basis, can be in fact supported by vastly different neural networks which can extend across multiple core memory structures. This idea is made explicit by plotting separately each lesion effect for each training condition (Fig. 4). The performances of the sham groups in the spatial and response memory tasks are shown in the bottom two rows; these are highly similar. Each column above indicates the individual contributions of DSL, HPC, and DSM in one training condition; these are highly distinct within and across lesion groups. Thus, behaviour is highly degenerate.

**Fig. 4.**
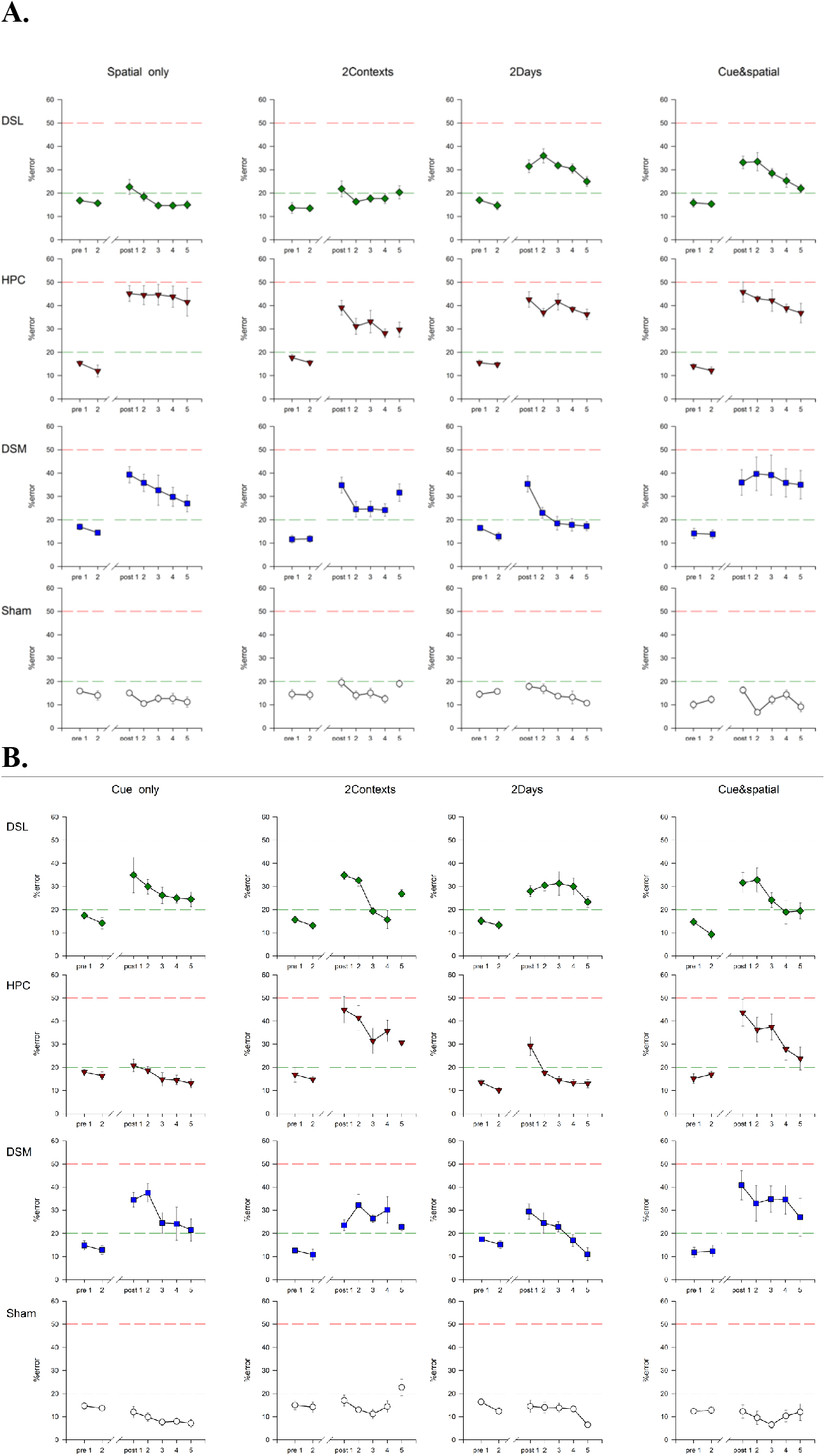
Normal behavior is highly degenerate. Same data as in Fig. 2C replotted to facilitate visualizing the distinct contributions DSL, HPC, and DSM had to normal behavior. The training protocol is indicated at the top of each column and the performance of the sham group in that condition is shown in the bottom panel. Comparing across the bottom row indicates that normal animals performed similarly regardless of training. However, in each case the performance was supported to different degrees by the three brain areas. **A. Spatial navigation. B. Cue response.**

## DISCUSSION

Patient H.M.’s selective amnesia that followed after the bilateral resection of his medial temporal lobes (Scoville & Milner, 1957) indicated that memory is not a general property of neural networks. Instead, the pattern of memory deficits documented in H. M. and similar patients showed that localized neural circuits are responsible for different kinds of memories. This discovery triggered numerous subsequent studies in which the mnemonic functions of various brain areas were extensively investigated. The analytical approach provided much information about memory organization in the brain and the underlying mechanisms. However, the opposite problem, of how different memories are integrated, remains poorly understood. The current data contribute to filling some of this gap, as they demonstrate that the DSL and HPC link different types of memories based on spatial and temporal factors, respectively. Notably, the memories involved were of distinct kinds, suggesting a real possibility that activity in one memory circuit may affect processes in a different memory circuit(Kim & Baxter, 2001). Despite the simplicity of the experimental manipulations, the implications of these results are far reaching and lead to significant questioning of our current perspective on memory architecture.

The ability of the HPC circuits to link different types of memories, primarily by using temporal rather than spatial factors, indicates that the ‘when’ of episodic memory may act as the thread linking the ‘where’ (and possibly also the ‘what’) of individual events, a prospect also suggested by recent data (Rubin, Geva, Sheintuch, & Ziv, 2015; Cai et al., 2016; Mankin, Diehl, Sparks, Leutgeb, & Leutgeb, 2015). Thus, the HPC is critical for spatial representations in general, but it can additionally support response memories if they are formed in close temporal proximity to spatial memories. This property of the HPC network, complementary to the DSL’s ability to link memories based on context, dovetails well with the quintessence of episodic memory. We recollect as a coherent unit events that occur in close succession regardless of environment, and distinguish among events that happen in the same space at different points in time. If cognitive memories can be thoroughly integrated with habits within the HPC, then it is not surprising that habits, which can be performed without conscious awareness (as for example in typing) can also be consciously recollected.

A second effect of a memory architecture in which HPC can be integral part of the neural network supporting habits is that a habit can be rescued after damage to the neural network primarily responsible for its acquisition (i.e., DSL). The current data supported this hypothesis. When animals acquired the response and spatial memories concurrently and HPC was involved in response memory, DSL lesions caused a transient rather than permanent deficit in response memory (Fig. 4B, second and fourth panels of first two rows). In contrast, when the response memory was formed either by itself or in temporal separation from the spatial memory, both cases in which the HPC network did not contribute to the behaviour, the impairment was long lasting (Fig. 4B, first and third panels of first two rows). This fact also means that the persistence of marked impairment in the response task after DSL lesions in the 2Days condition (Fig. 4B, third panel of the first row) does not reflect a ‘weaker’ response memory trace following training spaced across days. Thus, these data strongly support the idea that HPC can independently support S-R behaviour. The above analysis however should not be interpreted as implying that HPC and DSL contributions to behaviour combine in linear fashion.

The complementary aspect of DSL’s functional role revealed by the current data is that while DSL supports a rigid habitual response to a single cue in general, it can also become involved in a behaviour illustrative of cognitive memory if the two types of learning occur in the same space. This property of the DSL network may be involved in drug relapse, whereby exposure to an environment previously associated with drug abuse (an extreme form of habit) leads to renewed drug consumption. Even as the results of the reversal test in the 2Contexts condition suggested that context representations were selectively dependent on the HPC, reports that context memories can become HPC independent (Lehmann et al., 2009; Sparks, Lehmann, & Sutherland, 2011) and that the DSL can be involved in context memories (White & Salinas, 2003) leave open the possibility that along with the HPC, the DSL may also form its own context representation. The precise neural processes than enable DSL to support spatial and response memories acquired in the same environment remain currently unknown, although they may be related to the functional heterogeneity of this structure (Vicente, Galvão-Ferreira, Tecuapetla, & Costa, 2016) and to the dynamic reorganization of the DSM-DSL circuits during skill learning (Yin et al., 2009). Whatever contextual representation DSL may form, it is however insufficient to support spatial navigation on its own (Fig. 4A, last two panels of the first two rows), a result in agreement with the permanent spatial deficit seen in animals with HPC lesions that have undergone extensive spatial training (Clark, Broadbent, & Squire, 2005). The asymmetry between the capabilities of the two structures relative to behaviour may have an evolutionary basis: DSL and HPC evolved at different points in time during phylogeny and thus, the HPC built on an already existent DSL network (Murray, Wise, & Graham, 2016). Collectively, these findings have important implications for addiction. If the HPC network can independently support a habit while the DSL network can use context to link habits and cognitive memories, then treating addiction has to address simultaneously striatal and hippocampal processes from the perspective that these two networks operate as an integrated unit rather than as separate modules, each with a distinct contribution along a set function (cf. Ferbinteanu, 2019).

A final comment regards the DSM. As predicted by a role in behavioural flexibility, DSM supported both spatial and response memories regardless of training condition (presumably because both tasks require flexible switch between right and left body turns at the centre of the maze), and its lesion caused a somewhat larger memory deficit when the animals had to switch between tasks in the same training session (2Contexts condition). In the context of the current work, the effects of the DSM lesions further support the idea that the processes through which the DSL and HPC link distinct types of memories are not of a general nature but rather specific to each local network. That is, the distinct and complementary roles HPC and DSL play in integrating memories are not a result of processes that take place in memory circuits in general, but emerge from phenomena characteristic to each of these structure whose details will have to be elucidate in future work.

In conclusion, the current data indicate that by using time and space, HPC and DSL can integrate cognitive and habitual memories in distinct and complementary fashion. This finding provides an articulated framework within which to further investigate the underlying brain mechanisms that lead to the formation of coherent, multifaceted memories, and underscores the degenerate nature of behaviour. Together with previous research (Ferbinteanu, 2016), the current work helps qualify the multiple memory systems paradigm. Different types of memories are indeed critically dependent on distinct neural circuits. In some situations, these circuits operate independently and in parallel to guide behaviour. However, in some other situations, the neural basis of a given type of memory can be greatly expanded across multiple memory structures while at the same time basic features of experience, such as time and space, can be used to integrate different types of memories within the same local network.

## ACKNOWLEDGEMENTS

I thank Kasey Stern for her work on this project during her 2019 summer internship in my lab and Dr. Frank Barone for access to equipment in his lab. I would also like to acknowledge Drs. Robert McDonald, Bryan Devan, Mark Packard, Jarid Goodman, and Norman White for comments on previous versions of the manuscript. This work has been supported by NIH grant MH115421.

